# Herbarium specimens as tools for exploring the evolution of biosynthetic pathways to fatty acid-derived natural products in plants

**DOI:** 10.1101/2023.05.08.539892

**Authors:** Emma Fitzgibbons, Jacob Lastovich, Samuel Scott, Amanda L. Grusz, Lucas Busta

**Affiliations:** Department of Chemistry and Biochemistry, University of Minnesota Duluth, Duluth, MN, USA; Department of Biology, University of Minnesota Duluth, Duluth, MN, USA; Department of Botany, National Museum of History, Smithsonian Institution, Washington, DC, USA

**Keywords:** plant chemistry, metabolic evolution, herbaria

## Abstract

Plants synthesize natural products via lineage-specific offshoots of their core metabolic pathways, including fatty acid synthesis. Recent studies have shed light on new fatty acid-derived natural products and their biosynthetic pathways in disparate plant species. Inspired by this progress, we set out to expand the tools available for exploring the evolution of biosynthetic pathways to fatty-acid derived products. We sampled representative species from all major clades of euphyllophytes, including ferns, gymnosperms, and angiosperms (monocots and eudicots), and we show that quantitative profiles of fatty-acid derived surface waxes from preserved plant specimens are consistent with those obtained from freshly collected tissue. We then sampled herbarium specimens representing >50 monocot species to assess the phylogenetic distribution and infer the evolutionary origins of two fatty acid-derived natural products found in that clade: beta-diketones and alkyl resorcinols. These chemical data, combined with analyses of 26 monocot genomes, suggest whole genome duplication as a likely mechanism by which both diketone and alkylresorcinol synthesis evolved from an ancestral alkylresorcinol synthase-like polyketide synthase. This work reinforces the widespread utility of herbarium specimens for studying leaf surface waxes (and possibly other chemical classes) and reveals the evolutionary origins of fatty acid-derived natural products within monocots.

**Significance Statement:** Plant chemicals are key components in our food and medicine, and advances in genomic technologies are accelerating plant chemical research. However, access to tissue from specific plant species can still be rate-limiting, especially for species that are difficult to cultivate, endangered, or inaccessible. Here, we demonstrate that herbarium specimens provide a semiquantitative proxy for the cuticular wax profiles of their fresh counterparts, thus reducing the need to collect fresh tissue for studies of wax chemicals and suggesting the same may also be true of other plant chemical classes. We also demonstrate the utility of combining herbarium-based plant chemical profiling with genomic analyses to understand the evolution of plant natural products.

## 1. Introduction

Plant chemicals are a cornerstone of medicine and are also of great importance in agriculture and biotechnology (1). Plant chemicals thus relate to several of the United Nations’ Sustainable Development Goals, particularly those aimed at zero hunger as well as good health and well-being, since an advanced understanding of plant chemistry can help minimize the negative environmental impacts of agriculture and ensure food security in the face of climate change. Our knowledge of plant chemistry has advanced considerably in past decades and has recently been accelerated by advances in nucleotide sequencing technologies, leading to major advances in our understanding of specialized or lineage-specific metabolic pathways (2). However, as studies of plant chemistry become wider in scope, they remain limited by a lack of access to tissue from diverse plant lineages, and researchers must spend considerable time, energy, and resources to obtain plant tissue for analyses. These difficulties are particularly pronounced when studying plants that are endangered, that grow in remote or difficult to access geographic areas, or that are challenging to cultivate. Until we identify straight-forward avenues by which to systematically access tissues from diverse plant species, our ability to advance fundamental knowledge of plant chemistry will be delayed. This, in turn, impedes the deployment of plant chemistry-based solutions for tackling climate change.

Outside of wild populations, much of global plant diversity is preserved withing systematized collections, including germplasm collections (seed banks), living collections (botanical gardens), and museum collections (herbaria). In recent decades, the plant voucher collections stored in herbaria have been transformed by digitization, allowing for systematic evaluation of collections spanning hundreds of millions of specimens collected around the globe over centuries (3, 4). Not only is this wealth of preserved plant specimens an incredible source of data for biodiversity, morphological, and phenological studies (5, 6), it can also provide access to diverse taxa for chemical sampling of, for example, terpenoids, flavonoids, alkaloids, and glucosinolates (7–11). In the context of plant chemical research, we propose that using herbarium specimens could be especially useful for studies of metabolic evolution, for identifying lineages with variant metabolic machinery, as well as for discovering structural variants of known natural products and new natural products alike. Thus, herbaria have the potential to be a source of material for diverse studies that advance our understanding of plant chemistry.

The goal of this study was to evaluate the potential of herbarium specimens in studies of plant chemical diversity and plant metabolic evolution. To meet this goal, we chose to study fatty acid-derived natural products, a diverse group of compounds with both structural and bioactive properties (12). These compounds can be found in cuticular waxes—mixtures of hydrophobic biochemicals that coat plant surfaces. We supposed that fatty acid-derived natural products could be extracted from the surfaces of small tissue fragments from herbarium specimens and then analyzed with gas chromatography-mass spectrometry as is currently performed with fresh tissue (13–15). In this report, we demonstrate that the wax chemical profiles obtained from preserved tissue (herbarium specimens) are reasonable proxies for chemical profiles resulting from analyses of fresh tissue. Then, we used herbarium specimens to screen a wide range of monocot species for two classes of fatty acid-derived natural products, alkyl resorcinols and beta-diketones. Finally, we combined the chemical profiles from herbarium specimens with existing reference genome data for monocot species to provide strong evidence for the relative evolutionary origins of these two compounds, as well as a genetic mechanism by which their biosynthesis genes may have evolved from an alkyl resorcinol synthase-like ancestor. Based on this success, we suggest that phytochemists further develop partnerships with curators of natural history collections to advance large-scale studies of phytochemistry and to hasten the implementation of plant chemistry-based responses to sustainable development goals.

## 2. Results and Discussion

The objective of this study was to explore the use of herbarium specimens in studying the diversity and evolution of fatty acid-derived natural products in plant wax extracts. To meet this objective, three major activities were undertaken: (i) waxes from corresponding fresh and preserved specimens were compared across representatives of eudicot, monocot, gymnosperm, and fern lineages, (ii) preserved specimens were used to explore the distribution of alkyl resorcinols and beta-diketones, two fatty acid-derived natural products, across the monocot lineage, and (iii) public sequence data were used to assess the evolution of alkyl resorcinols and beta-diketone synthases across the monocot phylogeny.

### A. Waxes from preserved specimens are highly similar to those from corresponding fresh tissue

An initial screen revealed striking similarities between the gas chromatography-mass spectrometry total ion chromatograms obtained from wax extracts from fresh and preserved tissues of *Turritis glabra* L. (Brassicaceae; Figure 1A). This suggested that information about compound occurrence, at least for some wax compounds, could potentially be obtained from analyses of preserved plant tissue. To evaluate this potential, our first objective was to perform a more thorough comparison of waxes from fresh and preserved specimens. We selected three species each from eudicot, monocot, gymnosperm, and fern lineages (total 12 species; Figure 1B) and obtained fresh and preserved leaf tissue samples for three biologically independent individuals from each of these species (total 72 tissue samples; where biologically independent samples were not available, distinct leaves were used instead). Cuticular waxes were extracted from each tissue sample using chloroform and the extracts were analyzed with gas chromatography-mass spectrometry. Among the sample analyzed, we identified wax compounds belonging to eleven different wax chemical classes. Some of these chemical classes were ubiquitous classes that have been found, in some amount, on the surfaces of virtually all plant species, including very-long-chain *n*-alcohols, *n*-aldehydes, *n*-alkanes, alkyl esters, and fatty acids. We also identified wax compounds that are found only in certain lineages, including alkyl resorcinols, as well as very-long-chain secondary alcohols, diols, ketones, monoacyl glycerides, and triterpenoids. Overall, the data suggest that preserved plant specimens could be a source of diverse wax compound classes. Previous comparisons of volatile terpenoids from herbarium specimens and fresh tissues revealed that herbarium specimens had a similar distribution of terpenoid compounds (terpenoid “composition”), but not necessarily similar total terpenoid amounts (9). It seems likely that surface waxes would behave similarly, given that some species waxes accumulate in thick surface layers (epicuticular wax crystals) that can be physically removed with even a gentle touch, let alone actions associated with collection, pressing, mounting, and storing. For that reason, the remainder of the analyses presented here focus on wax composition, not total wax amounts.

**Fig. 1.**
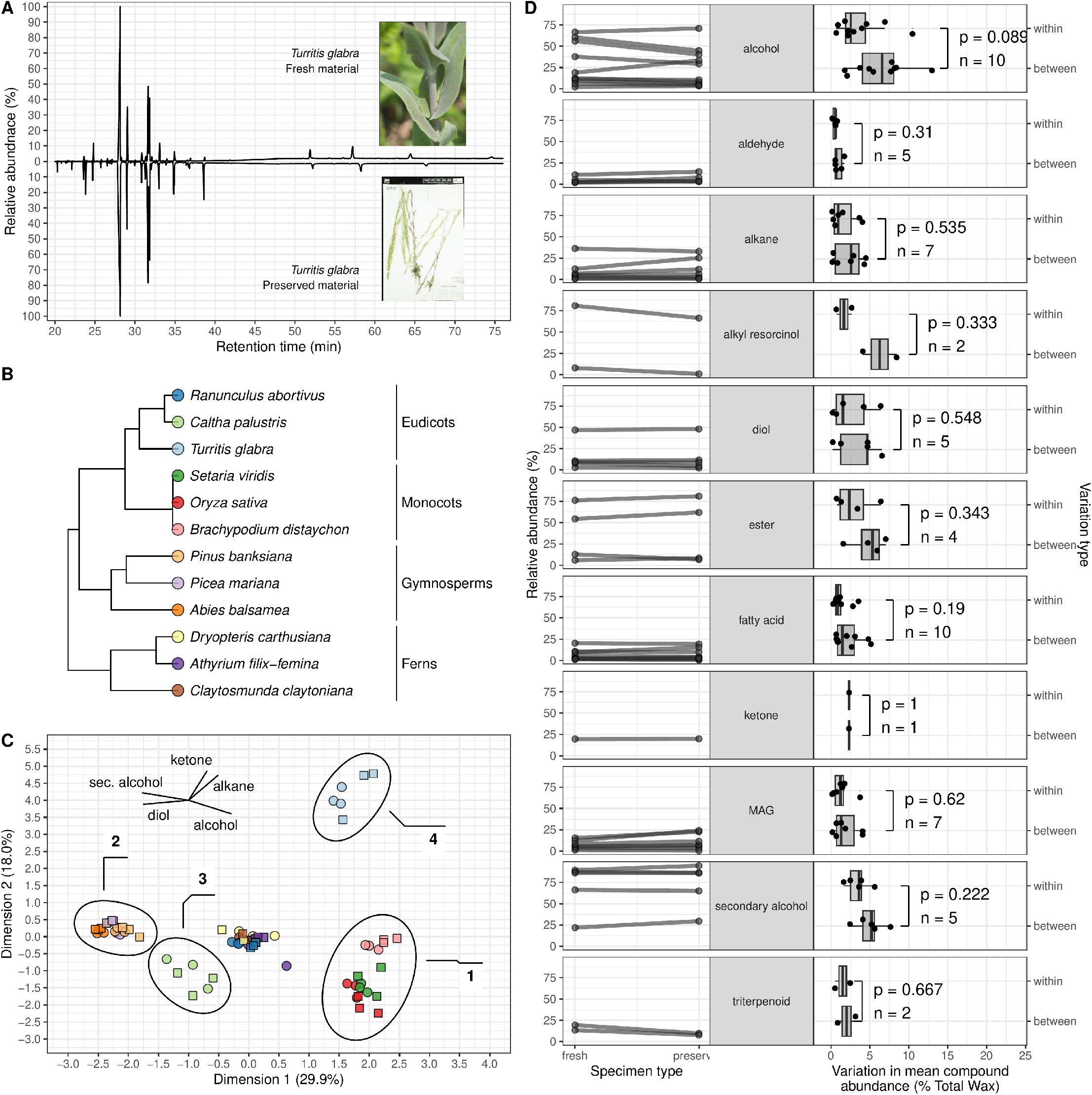
Comparison of waxes from fresh and preserved leaf tissue. **A**. Gas chromatography-mass spectrometry total ion chromatograms from analyses of cuticular waxes from fresh and preserved *Turritis glabra* tissue. The retention times of some late-eluting analytes do not exactly match because the samples were run under gas chromatography conditions with slightly different column lengths, however, the mass spectra for all corresponding compounds between the two samples were identical. **B**. Phylogeny of the 12 species selected for analysis. **C**. Scatter plot showing a principal components analysis of the 72 wax samples analyzed. Circles correspond to fresh tissue, squares to preserved tissue, with colors that correspond to the species-based color scheme in B. Black circles indicate the major distinguishing chemotypes identified among the species analyzed. The line segments in the upper left-hand corner are an ordination plot illustrating the compound classes that are most strongly aligned with the principal components. **D**. Quantitative comparison of the relative abundance of wax compound classes on fresh and preserved plant material. Line diagrams on the left connect the relative abundance of each compound class in corresponding fresh and preserved tissues. Slopes of the lines indicate the degree of difference in the relative abundance of a particular compound class between corresponding fresh and preserved specimens. The number of lines shown for each compound class indicates the number of species in which that compound class was found. The panels in the middle show the general structure of each compound class. Box plots on the right show the amount of variation in mean compound class relative abundance within and between a set of independent samples.

To evaluate the potential of waxes from preserved plant specimens to serve as quantitative proxies for extracts from freshly collected tissue, quantitative chemical profiles from the fresh and preserved tissue of 12 representative plant species were compared. Since different wax compound classes can degrade at different rates (i.e., different compound classes might be preserved to different degrees), this analysis was conducted on a compound class-by-compound class basis for each species. To visualize such trends in preservation, a plot was created in which the mean relative abundance of each compound class in fresh tissue was connected with a line to the mean relative abundance of that class in preserved tissue from that species (Figure 1D). The lines in the plot were relatively flat, indicating that the relative abundance of each compound class detected was highly similar in extracts of corresponding fresh and preserved specimens, and suggested that a statistical analysis should be devised to test for differences between these relative abundance values. However, it is well established that cuticular wax chemistry can change quantitatively in response to environmental stimuli (16–18), and it seemed highly likely that such variation might preclude direct quantitative comparisons of preserved and fresh tissue chemistry. Accordingly, in order to determine whether wax compounds differed in their quantitative contribution to overall wax mixtures, it was not the mean abundances between fresh and preserved tissue that were compared, but rather the variance in the abundance of each compound class between replicates of a given tissue type (fresh vs. preserved) versus the variance between fresh and preserved tissue. Under this scheme, if a compound class regularly degraded in a quantifiable way, the variance between fresh and preserved would be significantly greater than the variance within a tissue type. For the compound classes tested here, we compared these variances and found no significant difference for any compound class (Wilcox tests, α= 0.05, Figure 1D). This absence of differences suggested that each of these compound classes could indeed be preserved at a quantitative level. We acknowledge that such preservation may not be the case for all species or compound classes. For example, the inadvertent removal of epicuticular wax enriched in a given compound class would almost certainly distort correspondence between fresh and preserved tissue. However, the data presented here suggest that herbarium specimens are a suitable first approximation for the quantitative wax compositions found on fresh plant leaf tissue. It seems likely that this approach will also work for other classes of surface lipids, notably cutin, a polymer whose diversity is little studied compared with waxes.

### B. Based on preserved specimen chemistry, alkyl resorcinols are more widely distributed in monocots than are beta-diketones

Having identified that wax extracts from preserved specimens are reasonable proxies for extracts from corresponding fresh tissue, our next objective was to use herbarium specimen wax extracts to study the evolution of fatty acid-derived natural products. In particular, we targeted alkyl resorcinols and beta-diketones, two compound classes that have been reported from monocot lineages (19–21), but whose evolution has not been studied in detail. We began by examining wax extracts from preserved tissues of species previously reported to produce alkyl resorcinols or beta-diketones. Alkyl resorcinols, as trimethylsilyl derivatives, have a prominent peak in their mass spectra at m/z 268 (Figure 2A), so single ion chromatograms were extracted for that mass-to-charge ratio to search for alkyl resorcinols. In a wax extract from preserved tissues of *Trichophorum cespitosum* (L.) Hartm., which was previously reported to contain alkyl resorcinols (22), alkyl resorcinols were indeed identified as a homologous series (Figure 2B). Beta-diketones, as trimethylsilyl derivatives, have prominent ions in their mass spectra corresponding to alpha-cleavages adjacent to their secondary functional groups (Figure 2C). Accordingly, to search for diketones, gas chromatography runs were conducted in scan mode and then ion chromatograms were extracted for the ions that would be generated by a variety of possible beta-diketone structures including most common beta-diketone structures from grasses (Figure S1). For example, hentriacontan-14,16-dione was identified by its prominent mass peaks at *m/z* 325 and *m/z* 353 in preserved tissues of *Hordeum jubatum* L., a species previously reported to contain diketones (23). Thus, both alkyl resorcinols and beta-diketones can be detected in wax extracts from preserved plant tissues, and this process is facilitated with the use of extracted ion chromatograms.

**Fig. 2.**
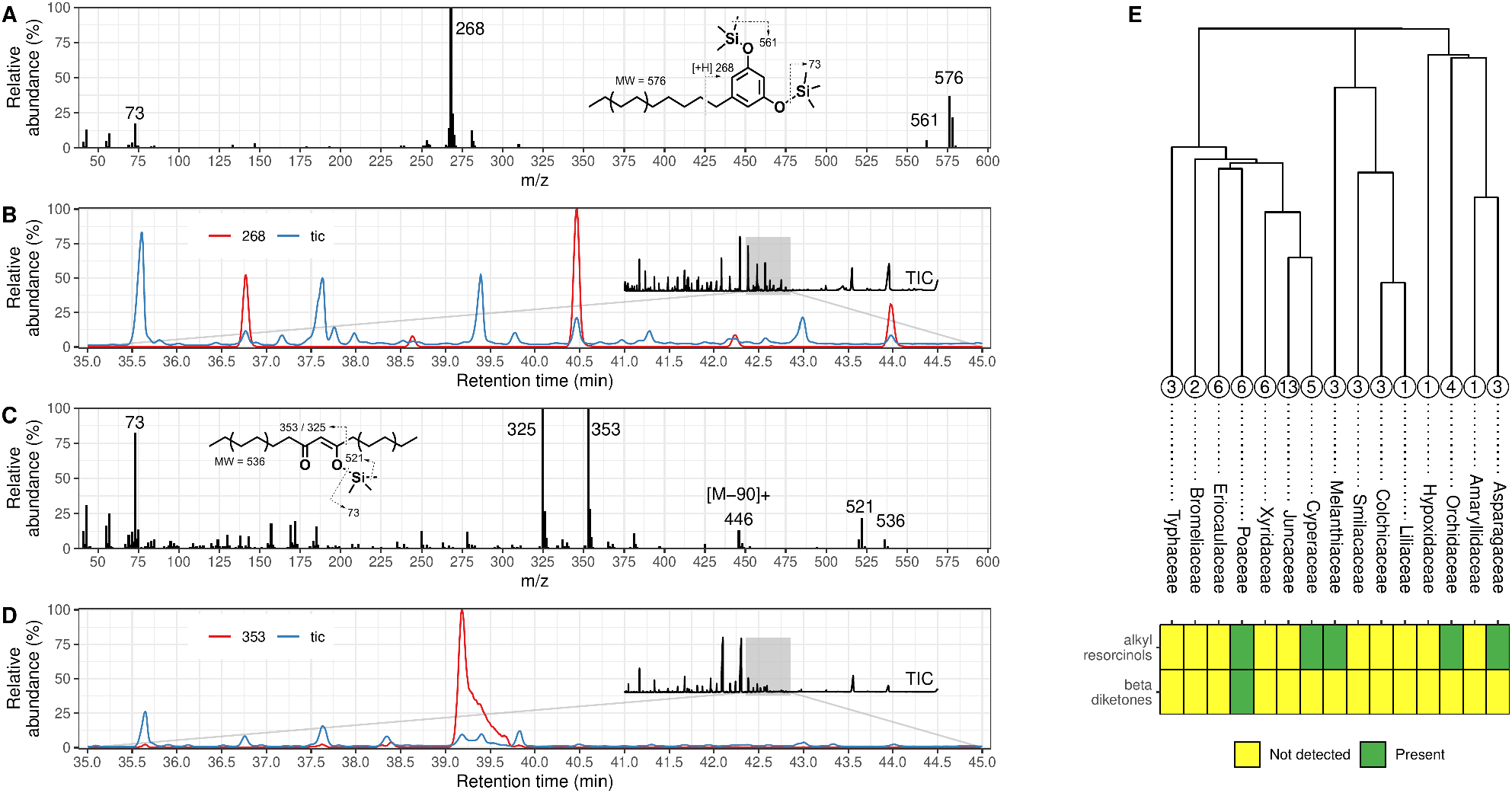
Occurrence of alkyl resorcinol and beta-diketone fatty acid natural products in monocot lineages. **A**. Mass spectrum of an alkyl resorcinol, with a prominent mass peak at *m/z* 268. **B**. Total and extracted (*m/z* 268) ion chromatograms of wax extracts from preserved tissues of *Trichophorum cespitosum*. The full total ion chromatogram is inset. **C**. Mass spectrum of a beta-diketone, with prominent mass peaks at *m/z* 325 and *m/z* 353. **D**. Total and extracted (*m/z* 353) ion chromatograms of wax extracts from preserved tissues of *Hordeum jubatum*. The full total ion chromatogram is inset. **E** Phylogeny of fifteen monocot families from which wax extracts from preserved tissues were analyzed. The numbers in the circles above each family indicate the number of species in that family that were analyzed. The heat map below the family names indicates the families in which alkyl resorcinols and beta-diketones were found, with green squares indicating that the compound class was found in that lineages and yellow indicating that the compound was not detected in that lineage.

Having confirmed that alkyl resorcinols and beta-diketones can be detected in wax extracts from preserved plant tissues, our next objective was to determine their relative times of evolution by determining whether alkyl resorcinols or betadiketones are more widely distributed across monocots. To meet that goal, wax extracts were prepared from preserved tissue of more than fifty monocot herbarium specimens spanning fifteen monocot families (Figure 2E). Across these samples, alkyl resorcinols were detected in at least one species from each of the families Poaceae (3 out of 6 species), Cyperaceae (5/5), Melanthiaceae (1/3), Orchidaceae (3/4), and Asparagaceae (2/3). In contrast, beta-diketones were only found on species within the Poaceae (1 out of 6 species). This suggests that alkyl resorcinols are more widespread in monocots than are beta-diketones and that the ability to synthesize alkyl resorcinols evolved before the ability to synthesize beta-diketones in this lineage.

### C. A phylogeny of monocot polyketide synthases suggests genome duplication facilitated diversification of ancestral alkylresorcinol synthase-like genes and the evolution of diketone synthesis

So far, the herbarium specimen-derived chemical data suggested that alkyl resorcinol synthesis predated beta-diketone biosynthesis in monocots. In addition, previous studies had identified alkyl resorcinol and beta-diketone synthesis genes in monocots (19, 24, 25), all of which are type-III polyketide synthases. Accordingly, our next objective was to use these characterized sequences and the previously assembled reference genomes of 26 monocot species (Table S3) to place our chemical data into a genomic context and to further explore the evolutionary origins of these two fatty acid-derived natural products. First, we used the amino acid sequences of the characterized alkyl resorcinol and beta-diketone synthases as queries for a BLAST and obtained a list of type-III polyketide synthases from the 26 monocot genomes. We filtered out low quality hits as well as hits that were clearly annotated as something other than a polyketide synthase or as a 3-ketoacyl CoA synthase. The sequences of the remaining hits were aligned, and the alignment was trimmed to remove positions with a high proportion of gaps (more than 70% gaps) or positions that had very low conservation (less than 30% conserved). From the trimmed alignment, we inferred a maximum likelihood phylogeny (Figure 3A). The overall topology of our phylogeny was highly similar to the monocot region of a previously reported type-III polyketide synthase phylogeny (26). In our phylogeny, the previously characterized monocot alkyl resorcinol and beta-diketone synthases were located in four large clades made up of sequences from the Poaceae (Clades 2A/2B and 3A/3B, Figure 3). These clades diversified after previously reported whole genome duplications in this lineage (nodes labeled “1”, “2”, and “3”; Figure 3; (26)). Gene duplications, from the single gene to whole genome level, have previously been associated with the functional diversification of genes involved in lineage-specific metabolism (2, 27–29). This precedent, along with the characterized alkyl resorcinol and beta-diketone synthases arising after the whole genome duplications, suggests that whole genome duplication played a major role in the functional diversification of type-III polyketide synthases in monocots.

**Fig. 3.**
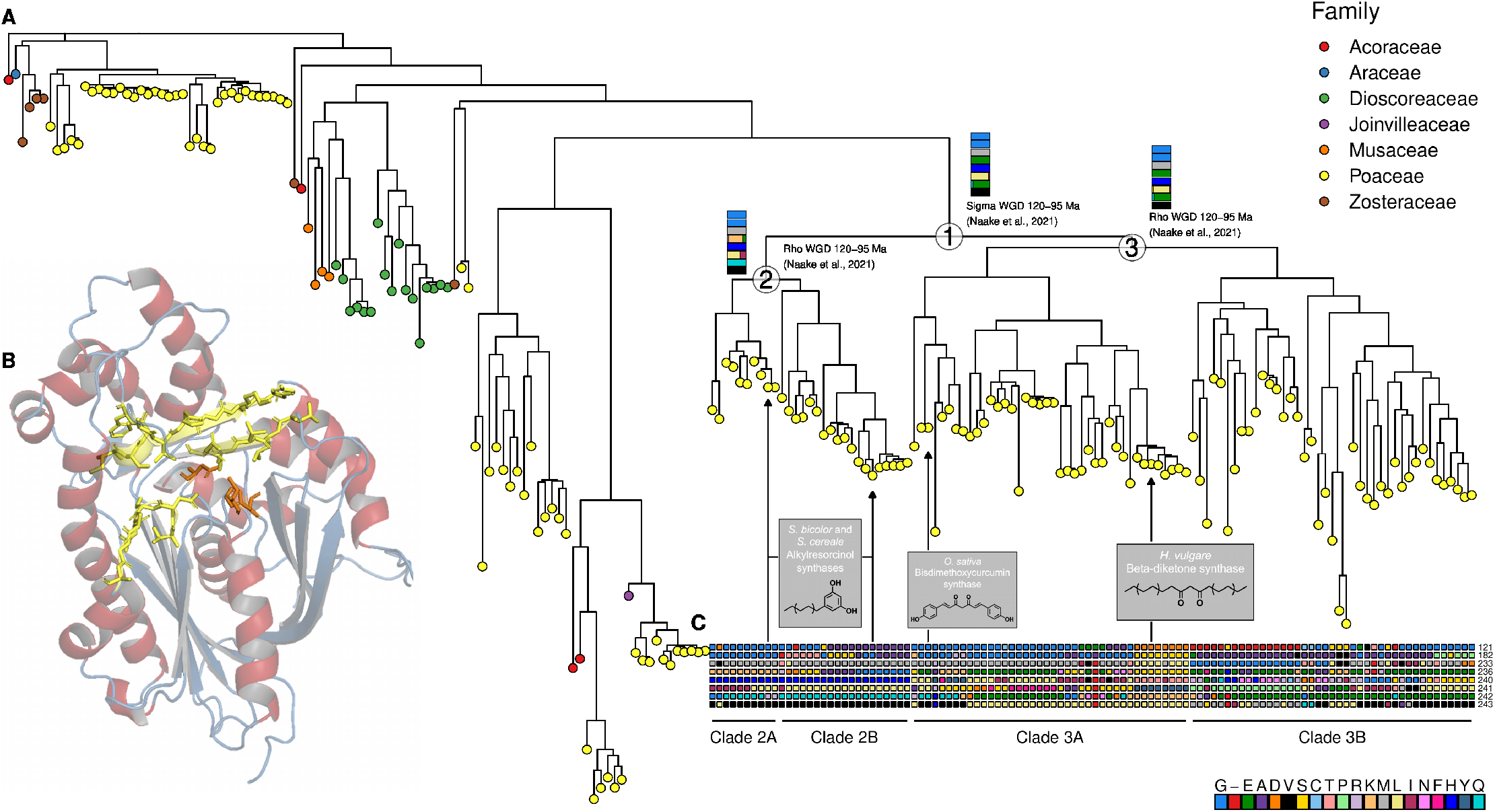
Characteristics of extant and ancestral polyketide synthases. **A**. Maximum likelihood tree of type-III polyketide synthase amino acid sequences from 26 monocot genomes. Tip color indicates the family in which the gene was found, as shown in the legend. The nodes labelled with numbered circles “1”, “2”, and “3” indicate whole genome duplications, with whole genome duplication at “1” preceding the divergence of clades 2 and 3 and whole genome duplications at “2” and “3” giving rise to 2A/2B and 3A/3B. **B**. A computer-generated model of a Poaceae type-III polyketide synthase basd on structurally similar crystal structures. The three orange residues are catalytic residues, and the eight yellow residues indicate amino acids near the active site pocket. **C**. Tiles indicating the identity of the eight active site amino acids in each of the proteins underneath the node labelled “1” in the phylogeny. Numbers on the right-hand side of the tiles indicate the position in our alignment in which each of these residues resides. The colors correspond to a particular amino acid as indicated in the legend. The horizontal bar plots at each whole genome duplication node in the phylogeny indicate the probability of the eight amino acid residues near the active site pocket in the ancestral sequences predicted for that node.

The finding that characterized alkyl resorcinol synthases and a characterized beta-diketone synthase arose from a common polyketide synthase ancestor via whole genome duplication raised questions about the catalytic abilities of the common ancestor. To learn more about this ancestor, as well as other ancestral polyketide synthase sequences, we first built a structural model of a plant type-III polyketide synthase and used it to identify positions in our type-III polyketide synthase amino acid alignment that correspond to active site residues (Figure 3B). We then mapped the identity of those active site residues onto the phylogeny (Figure 3C) and examined how those identities were distributed across the four clades arising after the two whole genome duplications. The two characterized alkyl resorcinol synthases and other members of both clades 2A and 2B had characteristic residues in certain position of the alignment (M233, H240, Q242, and V243; Figure 3C). In contrast, the characterized beta-diketone synthase did not have any of these characteristic active site residues (it instead had L233, N240, K242, and L243), something it shared in common with its closest neighbors in our phylogeny— genes from *Thinopyrum intermedium* (Host) Barkworth & D.R. Dewey, suggesting that species as a beta-diketone producer and one or more of those genes as beta-diketone synthases. Thus, analysis of alkyl resorcinol and beta-diketone synthase amino acid sequences revealed that the identities of the amino acids surrounding their active sites have diverged considerably.

Since eight active site residues so clearly distinguished alkyl resorcinol and beta-diketone synthases, we supposed that it might be possible to distinguish alkyl resorcinol synthase-like ancestral sequences from beta-diketone synthase-like sequences. Accordingly, we used maximum likelihood ancestral state reconstruction to estimate the identity of the same eight active site residues at the three nodes of the phylogeny that corresponded to whole genome duplications. At each of these nodes, the ancestral state reconstruction algorithm was able to estimate the identity of the eight residues with high certainty, and each predicted ancestor had characteristics that were strikingly similar to characterized alkyl resorcinol synthases (M233, H240, Q242, and V243) and absent from the beta-diketone synthase (Figure 3C, compare with colored bar charts at duplications). This finding strongly suggests that the ancestral polyketide synthase that gave rise to the sequences in clades 2A, 2B, 3A, and 3B (that is, both alkyl resorcinol synthases and the beta-diketone synthase) was very alkyl resorcinol synthase-like, if not an outright alkyl resorcinol synthase.

## 3. Conclusion

Here, we demonstrate that cuticular waxes from herbarium specimens are sources of fatty acid-derived natural products in proportions that are semiquantitatively consistent with the chemistry of fresh tissue. We then surveyed monocot wax chemicals and demonstrated that alkyl resorcinols are far more widespread across the monocot phylogeny than beta-diketones, which were only recovered from within Poaceae. We used ancestral state reconstruction and genomic analyses of monocot type-III polyketide synthases to reveal that, within the Poaceae, an ancestral alkyl resorcinol synthase-like gene appears to have diversified via whole genome duplication and given rise to beta-diketone synthases. Interestingly, those duplications have also generated several groups of polyketide synthases with virtually unexplored functions. It will be interesting to see what products’ synthesis they may encode. Overall, our findings suggest that herbarium specimens are a useful source of plant waxes for analytical study. Of particular fascination are the otherwise inaccessible gradients to which herbarium specimens may provide access (30–32), including analyses of waxes from plant tissues that were collected in the same location but separated by decades or even centuries of time.

## 4. Materials and Methods

### A. Plant Material and Chemical Analysis

Fresh plant material was collected from local green areas. Preserved plant samples were provided on loan by the Olga Lakela Herbarium (DUL) at the University of Minnesota-Duluth. To prepare a wax sample, a large hole punch was used to collect a disc of leaf tissue (2 cm in diameter) that was stored in a sealed scintillation vial. The discs of leaf tissue were extracted twice with 2 mL chloroform, the extracts were pooled, and the chrloroform was allowed to evaporate. The residue was then transferred to a GC vial insert, derivatized in 50 μl of 1:1 N,OBis(trimethylsilyl)trifluoroacetamide (BSTFA) and pyridine, and incubated (70 °C for 45 min.). Samples were analyzed on a 7890B Network GC (Agilent) equipped with an 7693A Autosampler (Agilent) equipped with a split/splitless injector and an HP-5 capillary column (Agilent, length 30 m x 0.250 mm x 0.25 μm film thickness); 2 μl of sample was injected on-column with He as the carrier gas with a flow rate of 1 mL/min. The initial temperature of the GC oven was 50°C and held for 2 min, followed by the first ramp at a rate of 40°C/min until it reached 200°C and was held for 2 min, then ramped at a rate of 3 °C/min until 320 °C and held for 30 min. The total run time for each sample was 77.75 min. with a solvent delay of 8 min. The analytes were detected using an Agilent 5977B (GC/MSD) Mass Selective Detector (EI 70 eV; m/z 40–800, 2 scans/s).

### B. Bioinformatics and Data Analysis

Species and family trees (Figure 1B and Figure 2E) were obtained by sampling from a published megaphylogeny (33). Principal components analysis was performed using the R package FactoMineR (34). BLAST searches, alignments, trimming, tree construction and visualization, and ancestral state reconstruction were performed as previously described (13) using the R packages “msa”, “gBlocks”, “phangorn”, and “ggtree” (35–38). Parameters for filtering BLAST hits were: length aligned with query > 250; hit length > 300 and < 600, e_value < 1E-50. Structural modeling was performed as previously described (39) using Phyre2 (40).

## 5. Acknowledgements

The authors wish to acknowledge support in the form of startup funds to LB from the Swenson College of Science and Engineering. We also extend our thanks to the University of Minnesota Duluth for supporting JL through the Undergraduate Research Opportunities Program. ALG is also grateful to NSF-DEB 2232106 for generous financial support. Specimens analyzed in this study were generously supplied by the Olga Lakela Herbarium (DUL).

## 7. Supplementary Information

- Supplemental Table 1: **Herbarium specimens used to assess wax chemical preservation**. Listed are the family, taxon, genus, species, replicate, and collector number. Entries marked with a dagger indicate instances in which biologically independent samples were not available and so independent leaves from the same individual were used instead.
- Supplemental Table 2: **Presence of beta-diketones and alkyl resorcinols in Poaceae**. Listed are the family, taxon, accession number for each specimen that was examined. The fourth and fifth columns indicate whether beta-diketones or alkyl resorcinols were found in that lineage.
- Supplemental Table 3: **Public sequence data used in this study**. Listed are the genus and species names of taxa whose genomes were used in this study. The data were obtained from Phytozome13 (https://phytozome-next.jgi.doe.gov/), with the file names listed alongside species names.

**Fig. S1.**
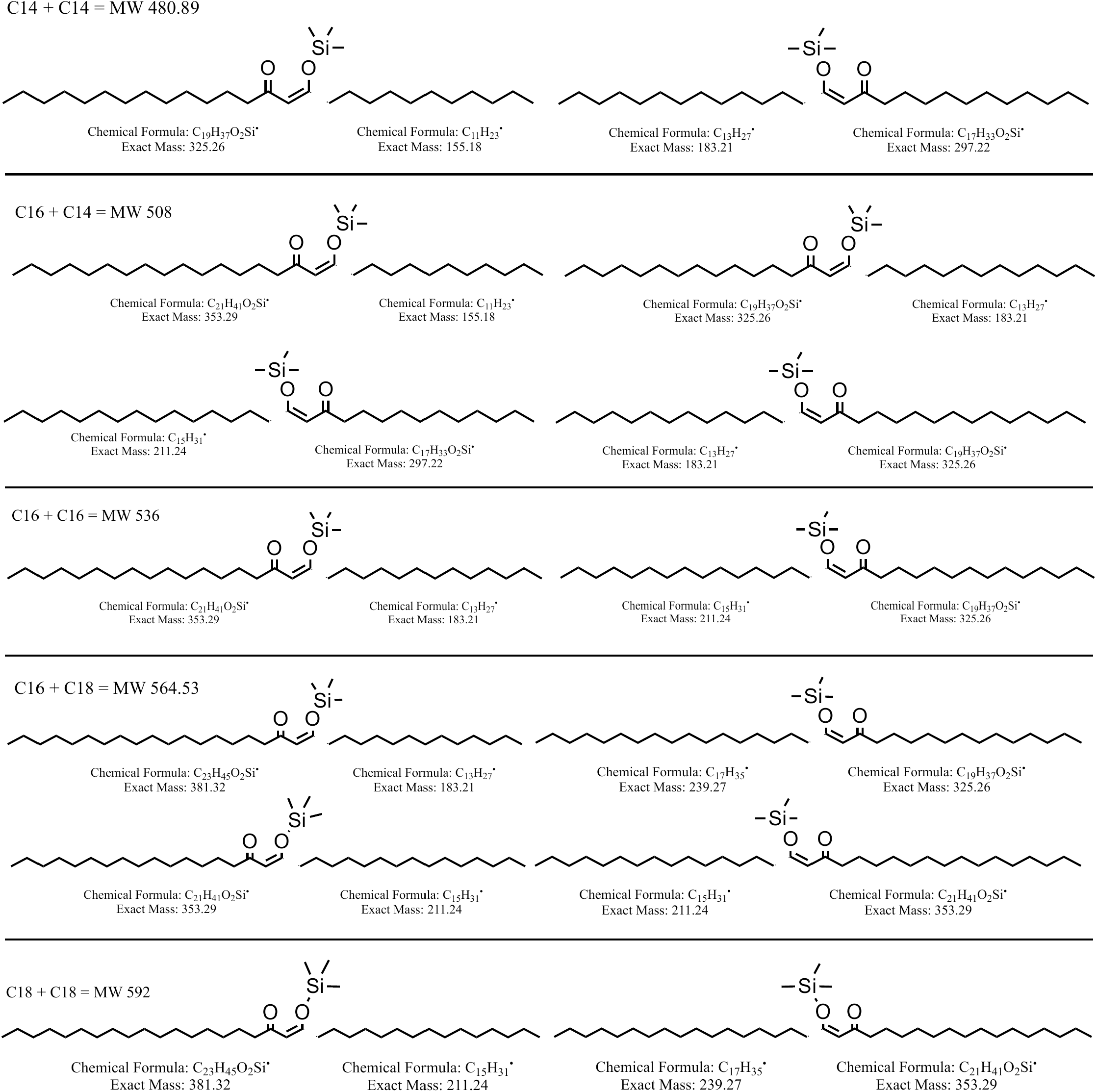
Fragmentation patterns of trimethylsilyl-derivatized beta-diketones. Each row shows structures for possible beta-diketones isomers and major fragments they may generate during mass spectrometry.

## Notes

### Competing Interest Statement

The authors have declared no competing interest.

